# Structure of human Mediator-RNA polymerase II transcription pre-initiation complex

**DOI:** 10.1101/2021.03.11.435010

**Authors:** Srinivasan Rengachari, Sandra Schilbach, Shintaro Aibara, Christian Dienemann, Patrick Cramer

## Abstract

Mediator is a conserved coactivator that enables regulated transcription initiation from eukaryotic protein-coding genes^1-3^. Mediator is recruited by transcriptional activators and binds the pre-initiation complex (PIC) to stimulate RNA polymerase II (Pol II) phosphorylation and promoter escape^1-6^. Here we prepare a 20-subunit recombinant human Mediator, reconstitute a 50-subunit Mediator-PIC complex, and resolve the complex structure by cryo-EM at an overall resolution of 4.5 Å. Mediator binds with its head module to the Pol II stalk and the general transcription factors TFIIB and TFIIE, resembling the Mediator-PIC interactions in the corresponding yeast complex^7-9^. One end of Mediator contains the metazoan-specific subunits MED27-MED30, which associate with exposed regions in MED14 and MED17 to form the proximal part of the tail module that binds activators. The opposite end of Mediator positions the flexibly linked CDK-activating kinase (CAK) of the general transcription factor TFIIH near the C-terminal repeat domain (CTD) of Pol II. The Mediator shoulder domain holds the CAK subunit CDK7, whereas the hook domain contacts a CDK7 element that flanks the kinase active site. The shoulder and hook reside in the Mediator head and middle modules, respectively, which can move relative to each other and may induce an active conformation of CDK7 to allosterically stimulate CTD phosphorylation and Pol II escape from the promoter.

---

Mediator consists of the head, middle and tail modules, which together comprise 21 and 26 subunits in yeast and human, respectively^1-3^. Mediator additionally associates with a 4-subunit kinase module that is however not part of the Mediator-PIC complex^10-12^. Structural studies of Mediator were long hampered by its large size and flexibility and concentrated on the yeast Mediator^3,13-15^. Detailed structures were obtained for the head module^16-18^ and a unified module architecture was derived^19-21^. The functional core of Mediator includes the head and middle modules^7,22^. The core Mediator structure highlighted an architectural role of subunit MED14^23^. The location of the yeast core Mediator on a PIC lacking TFIIH was determined and three Mediator-PIC interfaces were defined and named interface A, B, and C^7^. Structural analysis of a yeast Mediator-PIC complex confirmed the location of Mediator and revealed a contact between Mediator and a flexibly linked TFIIH module, the CDK activating kinase (CAK)^8^, which contains subunits CDK7, cyclin H, and MAT1^24^. The structure of a yeast core Mediator-PIC complex at 5.8 Å resolution confirmed the Mediator-CAK contact and provided more detailed insights into the Mediator-PIC interfaces^9^.

Despite this progress, the structures of human Mediator and its assembly with the PIC are lacking. It is thus unclear how human Mediator differs from its yeast counterpart and whether human Mediator interacts with the PIC in a conserved manner. Here we prepared a 20-subunit recombinant human Mediator that lacks only six subunits in the distal part of the tail module, and we determined the Mediator structure in complex with the PIC. The structure confirms that the Mediator core is conserved, reveals subunits in the proximal part of the tail module, and resembles the structure of murine Mediator that was described in a recent manuscript^25^. The human Mediator binds the PIC at the same position as its yeast counterpart, but there are also differences in the Mediator-PIC interfaces. We also define the contact between Mediator and the TFIIH CAK and uncover two interfaces with CDK7 that we call interfaces D and E.

## Recombinant human Mediator

Structural studies of human Mediator thus far relied on scarce and often heterogeneous endogenous preparations. To study human Mediator at high resolution, we produced recombinant Mediator after co-expression of its subunits in insect cells (Methods). Initial attempts to use 15 subunits that are orthologous to those present in the yeast core Mediator^23^ did not yield a well-defined preparation. However, when we additionally co-expressed the five subunits MED26-MED30, a soluble and pure 20-subunit recombinant human Mediator complex was obtained (Extended Data Fig. 1a). The recombinant human Mediator contains the complete architectural subunit MED14, the head and middle modules, and the proximal part of the tail module that binds the head^20^. The recombinant Mediator only lacks subunits MED1, MED15, MED16, MED23, MED24 and MED25, which form the distal part of the tail module that faces away from the PIC^8^. These efforts established a route to obtain milligram quantities of a homogeneous human Mediator for structural studies.

## Human Mediator-PIC structure

The PIC was prepared as described in the accompanying paper^26^ with recombinant human general transcription factors TBP, TFIIB, -E, -F, and -H^27^, and *Sus scrofa domesticus* Pol II^28^, which is 99.9% identical to human Pol II. The PIC was then incubated with recombinant human Mediator and ADP-BeF_3_, a transition state analogue for the catalytic TFIIH subunits XPB, XPD and CDK7 (Methods). The resulting Mediator-PIC complex (Extended Data Fig. 1b) was subjected to single-particle cryo-EM data collection (Extended Data Table 1). We then used 3D classification to identify a subset of particles containing the complete complex (Extended Data Fig. 2). Density for Mediator and TFIIH was improved with focussed classification and masked refinements. This led to a reconstruction of the Mediator-PIC complex at an overall resolution of 4.5 Å, although the focussed maps for Pol II and the Mediator head showed local resolutions of ∼3.5 Å and revealed side chain density (Extended Data Figs. 2-4).

To derive a model for the human Mediator-PIC complex, we placed the atomic PIC model in the closed promoter state that we derived in the accompanying study^26^, and fitted the RPB4-RPB7 stalk and parts of TFIIE into the cryo-EM density (Methods). The remaining unaccounted density was interpreted in four steps. First, the density for the Mediator head module was very well-defined and allowed for manual rebuilding of a homology model based on the yeast core Mediator structure^23^. Second, the Mediator middle module was more flexible, but the density showed secondary structure and enabled unambiguous rigid body fitting of a homology model for the middle module derived from the yeast core Mediator structure^23^. Third, unaccounted densities at one end of the core Mediator could be assigned to the Mediator subunits MED27-MED30 and modelled *de novo*. The last remaining density was unambiguously fitted with the known CDK7-cyclin H-MAT1 structure^29^. The resulting Mediator-PIC structure lacks only MED26 and shows good stereochemistry (Extended Data Table 1).

The overall human Mediator-PIC structure resembles its yeast counterpart, highlighting the high conservation of the general transcription initiation machinery throughout eukaryotes (Fig. 1). Human Mediator uses its head module to interact with the core PIC around the RPB4-RPB7 stalk of Pol II, occupying the same location on the PIC and adopting the same overall orientation as observed in the yeast Mediator-PIC complex^7-9^. Mediator binding does not alter PIC structure^26^, except for a change in the relative orientation of the RPB4-RPB7 stalk of Pol II and the TFIIE domain ‘E-ribbon’ that both interact with Mediator (Extended Data Fig. 5a). In the following, we will first describe the human Mediator structure and then discuss the Mediator-PIC interfaces and implications for Mediator function.

**Figure 1.**
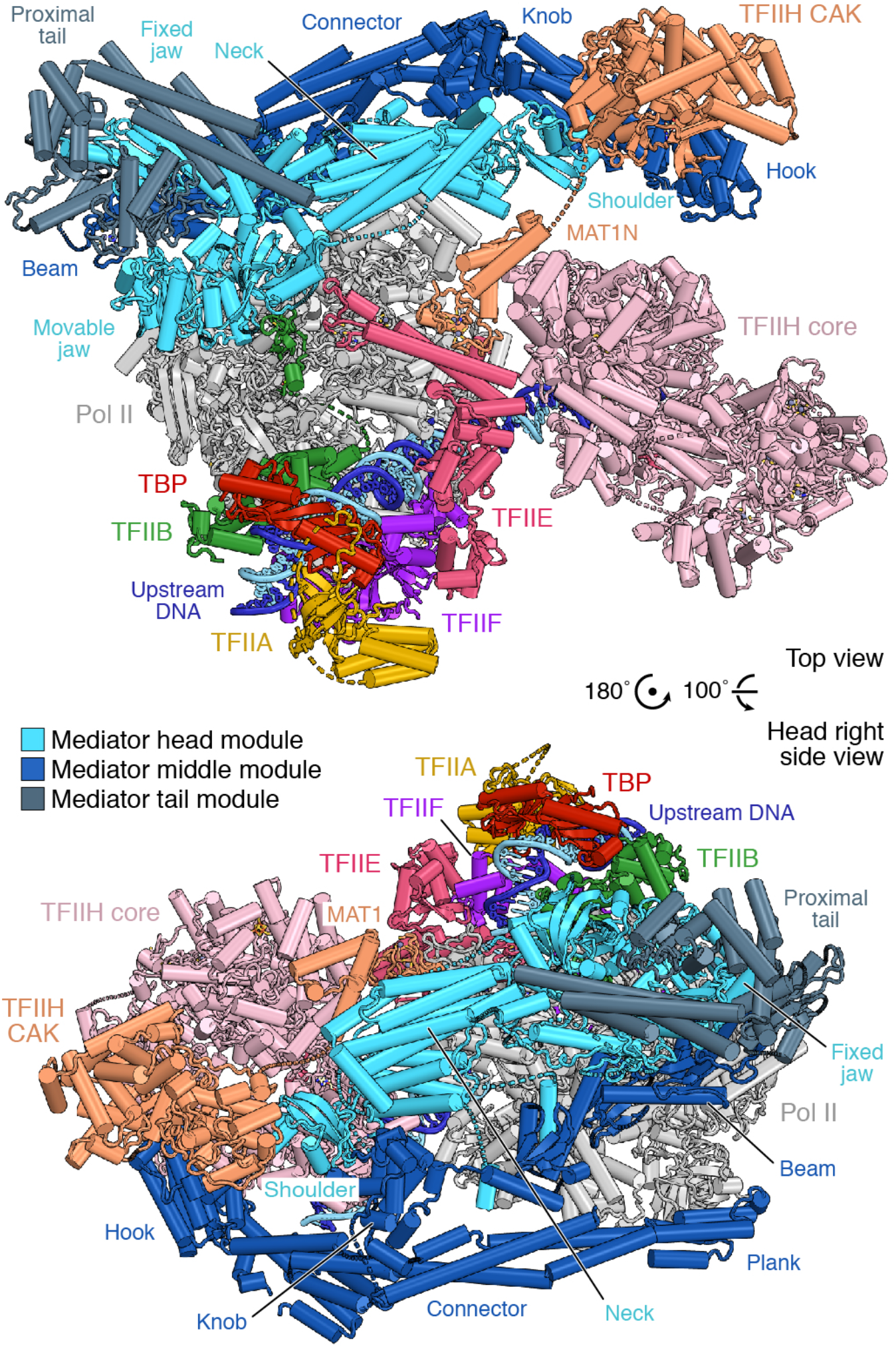
Human Mediator-PIC structure. Two views^9,36^ of the Mediator-PIC structure. TFIIH core and CAK modules are distinguished by colour. Positions of selected TFIIH subunits and Mediator domains within the head, middle and tail modules are indicated. The DNA template and non-template strands are in dark and light blue, respectively. Dashed lines represent flexible linkers. Colour code used throughout.

## Human Mediator structure

The structure of human Mediator resembles the previously defined core Mediator from yeast^23^, confirming the high conservation of Mediator throughout eukaryotes^30,31^ (Fig. 2, Extended Data Fig. 5b, Supplementary Fig. 1). In the head module, subunits MED6 and MED8 form the ‘neck’ region and the C-terminal helix of MED8, MED18 and MED20 form the ‘movable jaw’ (Fig. 2a). The head subunit MED17 shows some variations; its tooth domain is rotated and its nose domain is rotated and extended by an additional domain that we call ‘extended nose’ (Fig. 2b, c). The human Mediator also contains a helical insertion (residues 386-423) between its tooth and nose domains (Fig. 2d). The head subunits MED11 and MED22 contain longer C-terminal helices that protrude from the ‘fixed jaw’ of the head module (Fig. 2b, d). Comparison of our structure to the yeast core Mediator in its PIC-bound state^9^ reveals that the middle module is slightly rotated and shifted with respect to the head (Extended Data Fig. 5b). These movements are accommodated by ‘tethering’ helices that are flexibly linked to the head module subunits MED6 and MED17 and bind the middle module as observed for the yeast complex^9^. The ‘plank’ domain^7^ protrudes from the middle module and is mobile.

**Figure 2.**
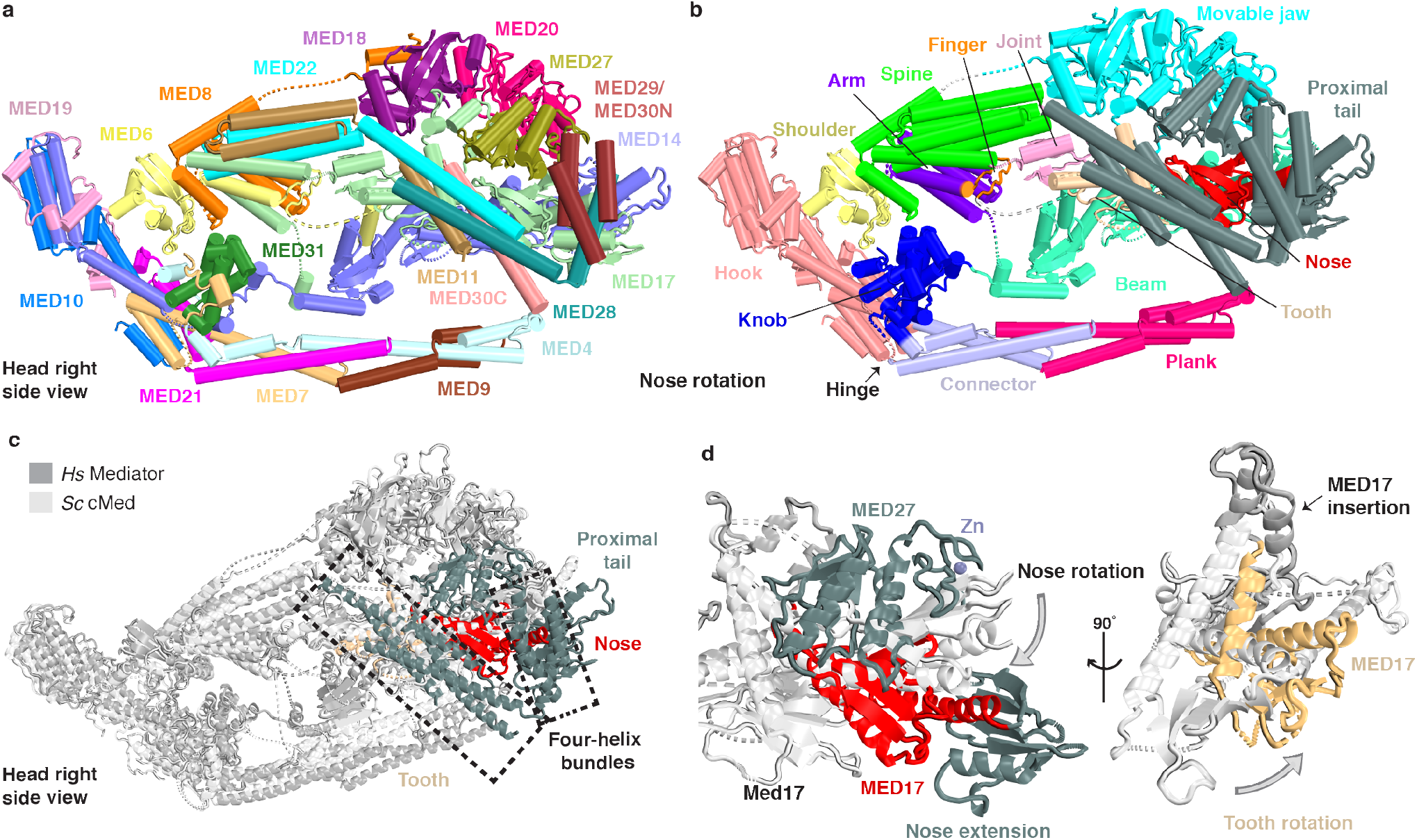
Features of human Mediator structure. **a.** Subunit architecture. Subunits are shown in different colors. **b.** Domain architecture. Domains are shown in different colors. **c.** Comparison with the yeast core Mediator structure (white). Altered MED17 regions and the proximal tail are highlighted. **d.** Alterations in human MED17 compared to its yeast counterpart (white).

Our structure additionally reveals the proximal part of the tail module (‘proximal tail’), which includes the newly observed subunits MED27-MED30 (Figs. 1, 2b). The proximal tail is anchored on the head module. MED27 forms a RWD domain with a C-terminal ‘zinc finger’ motif and nestles in between the fixed and the movable jaws of the head module. MED27 also contacts MED20, the extended C-terminal region of MED17, including the extended nose, and a region in the MED14 beam that alters its orientation. MED28 and MED30 contain long C-terminal helices that form a four-helix bundle with the C-terminal helices of MED11 and MED22. In MED28, a preceding helix is oriented at right angles to the C-terminal helix and is part of a second four-helix bundle that is poorly resolved due to its peripheral location. Our PIC-bound human Mediator structure is generally consistent with cryo-EM studies of the free murine Mediator^32^. Recently, a manuscript describing a structure of the murine Mediator became available^25^, but a detailed comparison awaits release of the coordinates.

## Mediator-core PIC interfaces

Our new structure shows that human Mediator binds the core PIC by formation of the previously described^7,9^ interfaces A and B (Fig. 3). In interface A, the movable jaw of the head module uses its MED18 subunit to contact the TFIIB B-ribbon and the TFIIE E-ribbon (Fig. 3b). Interface B is most extensive and is formed by interactions of the neck submodule in the Mediator head with the Pol II stalk (Fig. 3c). In particular, the arm domain of MED8 and the spine domain of MED22 both contact the stalk subunit RPB4, including its C-terminal helix (Fig. 3c). Compared to its yeast counterpart, the neck resides slightly closer to the Pol II stalk, resulting in more pronounced contacts with the core PIC. We do not observe the previously described^7,9^ interface C, formed between the Mediator plank and the Pol II foot domain, but movements in the middle module may enable formation of a similar contact.

**Figure 3.**
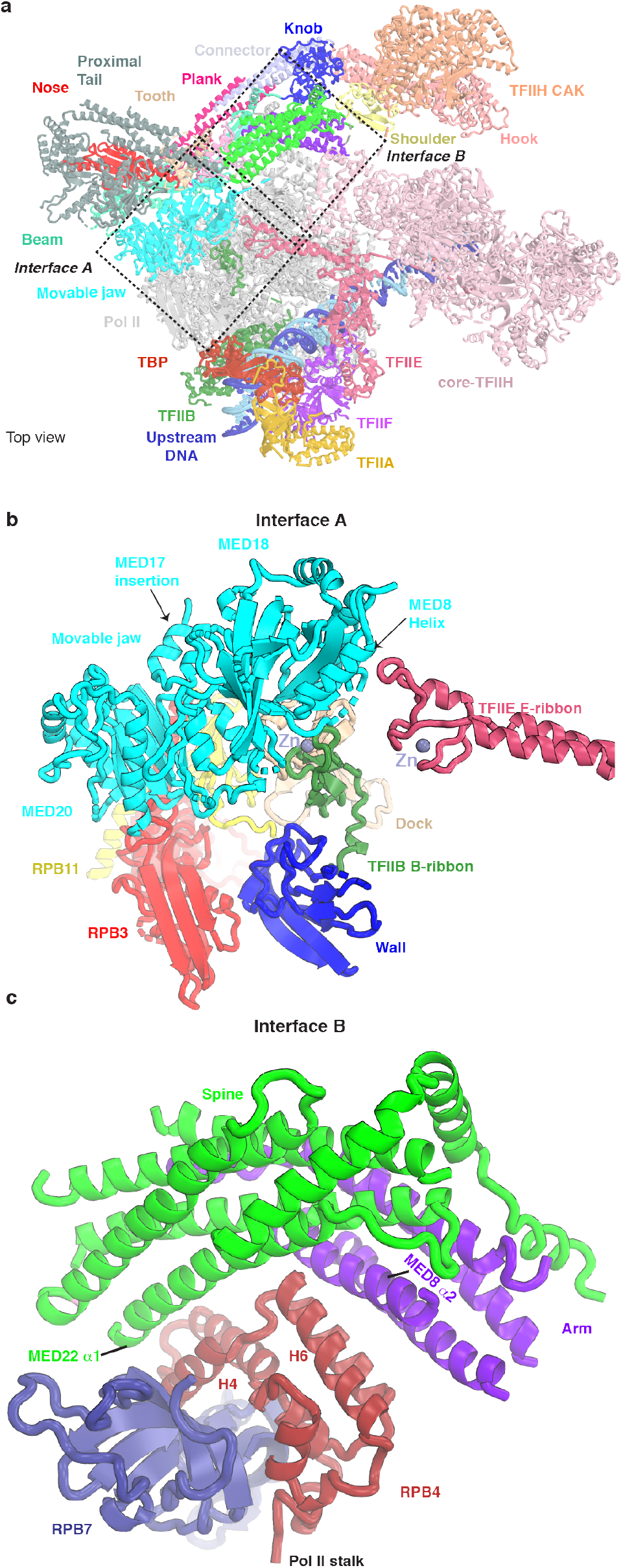
Mediator-core PIC interactions. **a.** Human Mediator-PIC structure with Mediator domains colored as in Fig. 2b. The location of the Mediator-PIC interfaces A and B are indicated. **b.** Interface A shows interactions between the Mediator head submodule ‘movable jaw’, the TFIIB B-ribbon and the TFIIE E-ribbon. **c.** Interface B shows interactions between the Mediator head submodule ‘neck’ and the Pol II RPB4-RPB7 stalk.

## Mediator-TFIIH kinase interactions

Our structure also reveals the TFIIH module CAK, which is anchored to the PIC via its subunit MAT1 (Fig. 4). Whereas the N-terminal region of MAT1 binds the TFIIH core and the Pol II clamp, the C-terminal region (MAT1C) associates with the CDK7-cyclin H pair. The CDK7-cyclin H-MAT1C trimer is flexibly linked to the rest of the PIC^26^, but in our structure it adopts a defined location that is similar to that in yeast Mediator-PIC complexes^8,9^. In contrast to previous studies, we can however define the orientation of the CDK7-cyclin H-MAT1C trimer and its interactions with Mediator.

**Figure 4.**
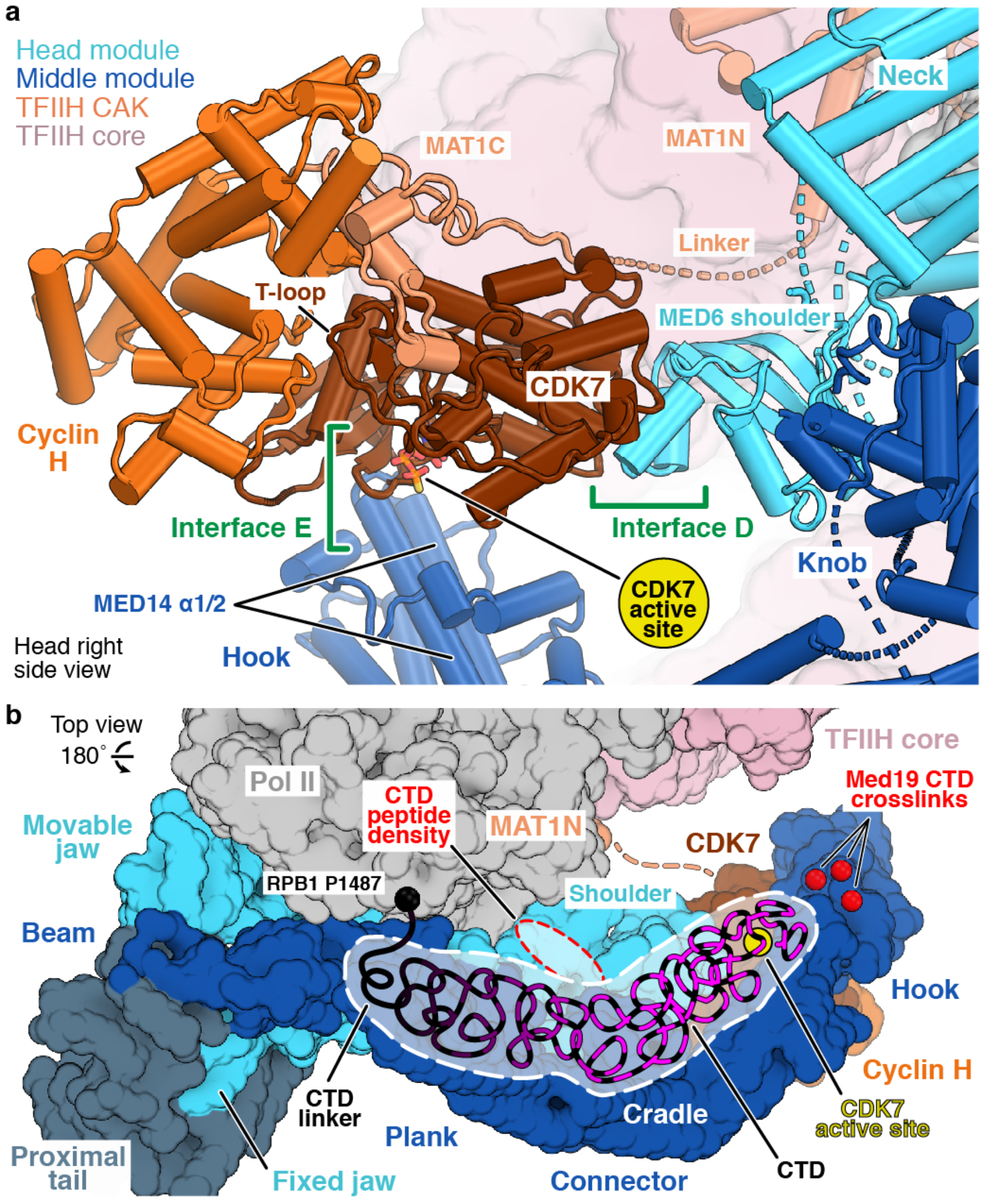
Mediator-TFIIH kinase interactions. **a.** Interfaces D and E. The location of the CDK7 active site is highlighted by modeling of an ATP analogue^29^. **b.** Putative path of the CTD from the CTD linker (black) through the cradle to the active site of CDK7 (yellow circle). Indicated are known sites of CTD crosslinks^9^ and density for a CTD peptide bound to the Mediator head^18^, both obtained from the yeast system.

Mediator contacts only the CDK7 subunit of the CAK and forms two distinct interfaces that we call interfaces D and E (Fig. 4a). Interface D is formed by contacts between the N-terminal region of the shoulder domain in the Mediator head subunit MED6 and two CDK7 regions that are specific to CDK7^33^ (residues ∼103-110 and ∼259-266). Interface E is formed between the MED14 N-terminal helices α1 and α2 in the hook domain of the Mediator middle module and the CDK7 N-terminal region (residues 10-26) that flanks the ATP-binding site and active center. In other CDKs, this N-terminal region harbours a ‘glycine rich loop’ that is involved in kinase activation^34^. Whereas interface D is more extended and may hold the CDK7-cyclin H-MAT1C trimer in position, interface E is likely transient because the Mediator hook is mobile.

These observations have implications for understanding how Mediator stimulates CDK7-dependent phosphorylation of the Pol II CTD. First, Mediator orders and positions the flexibly linked CDK7-cyclin H-MAT1C trimer. Second, Mediator orients the CDK7-cyclin H-MAT1C trimer such that the CDK7 active site faces the previously described cradle^7^, an open space that is formed between Mediator and the PIC. The orientation of CDK7 may facilitate targeting of the CTD substrate to the kinase, as the CTD could extend easily from the Pol II linker through the cradle to reach the CDK7 active site (Fig. 4b). This path is generally consistent with crosslinking sites^9^ and with density for a CTD peptide^18^ that were both observed in the yeast system. Third, the Mediator hook may stimulate kinase activity by allosterically inducing an active conformation of the CDK7 catalytic center. Whether the Mediator hook may cooperate with the CDK7 T-loop that is involved in kinase activation^29,35^ remains to be investigated.

## Conclusions

Together with the accompanying paper^26^, our work provides the structural basis for Mediator-regulated transcription initiation at eukaryotic protein-coding genes. Comparison with the previous yeast Mediator-PIC structure highlights the high conservation of the transcription initiation machinery throughout eukaryotes. Future work can now focus on different functional states of the Mediator-PIC complex and its possible alterations by activators. In addition, our recombinant human Mediator enables a dissection of the gene regulatory mechanism.

## Acknowledgements

We thank past and present members of the Cramer laboratory, particularly U. Neef, P. Rus and F. Grabbe for maintenance of the insect cells culture and purification of the recombinant human initiation factors. We thank C. Dienemann and U. Steuerwald for maintenance of the cryo-EM facility. S.R. was supported by a Peter-and-Traudl-Engelhorn postdoctoral fellowship. S.A. was supported by H2020 Marie Curie Individual Fellowship (894862). P.C. was supported by the Deutsche Forschungsgemeinschaft (EXC 2067/1 39072994, SFB860, SPP2191) and the ERC Advanced Investigator Grant CHROMATRANS (grant agreement No 882357).

## Author contributions

S.R. carried out experiments and data analysis, except for the following. S.S. established human PIC preparation. S.A. provided human PIC coordinates. S.A. and S.S. assisted with data processing and structural modelling. C.D. assisted with cryo-EM grid preparation and data collection. P.C. designed and supervised research. S.R., S.S., S.A. and P.C. interpreted the data and wrote the manuscript.

## Author information

Reprints and permissions information is available at www.nature.com/reprints. The author declare that they have no competing financial interest. Correspondence and request of materials should be addressed to P.C.

## Competing interests

The authors declare no competing interests.

## Data availability statement

The cryo-EM density reconstructions and final models were deposited with the EMDB (EMD-XXXX for Mediator + Pol II Stalk and EMD-XXX for Mediator + PIC) and with the PDB (YYYY). All data is available in the main text or the supplementary materials.

## METHODS

### Cloning and protein expression

Full length cDNA clones of 20 human Mediator subunits were either obtained from the Harvard Plasmid Repository or chemically synthesized by Integrated DNA Technologies (IDT). cDNAs were used as PCR templates for cloning into modified pFastBac vectors (derivatives of 438-A and 438-C; Addgene 55218 and 55220) by ligation independent cloning (LIC)^37^. All subunits were cloned without an affinity tag, except for a N-terminal His6-MBP-tag (MBP, maltose-binding protein) followed by a TEV protease site on Med17 and a N-terminal MBP-tag on Med14 with a cleavable TEV protease site. Three different constructs for baculovirus expression were generated, one containing the seven subunits of the head module (MED6, MED8, MED11, MED17, MED18, MED20, MED22), one for head-associated^20^ metazoan-specific subunits (MED26, MED27, MED28, MED29, MED30), and one for the middle module subunits (MED4, MED7, MED9, MED10, MED14, MED19, Med21, Med31). We refer to these three constructs as C1, C2 and C3, respectively. Whereas C1 and C2 were generated by biGBAC cloning methodology^38^, C3 was made using multiple rounds of LIC^37^. Final clones were verified by sequencing. Bacmid preparation and virus production (V0 and V1 stages) were performed as described^39^. Insect cell expression of Mediator was achieved by co-infection of the V1 virus for the constructs C1, C2 and C3 in Hi5 cells (Expression systems). After 60 hours of expression, cells were harvested by centrifugation (900 g, 10 min, 4 °C) and resuspended in buffer-A (25 mM 4-(2-hydroxyethyl)-1-piperazineethanesulfonic acid (HEPES) pH 7.5, 300 mM NaCl, 10% glycerol (v/v), 0.5 mM tris(2-carboxyethyl)phosphine (TCEP), 0.284 µg/ml leupeptin, 1.37 µg/ml pepstatin A, 0.17 mg/ml PMSF, 0.33 mg/ml benzamidine). The cell suspension was flash frozen in liquid nitrogen and stored at −80 °C.

### Protein purification

All steps of protein purification were performed at 4°C. Recombinant Mediator was purified by successive steps of affinity chromatography followed by ion exchange chromatography and size exclusion chromatography (SEC). Stored insect cell suspension was thawed at 25 °C. Cells were lysed by sonication and clarified by centrifugation (79,000 *g*; 60 min). The supernatant was passed over amylose resin (New England Biolabs) packed in an Econo-column (Bio-rad) pre-equilibriated with buffer-A and then washed with 20 column volumes (CV) of buffer-A. cMED was eluted with buffer-B (25 mM HEPES pH 7.5, 100 mM NaCl, 10% glycerol (v/v), 0.5 mM TCEP) containing 100 mM maltose and incubated overnight with TEV protease. The cleaved MBP tag and TEV protease were removed by anion exchange chromatography using a Mono Q 5/50GL (GE Healthcare) column pre-equilibrated with buffer-B. Post sample application, the column was washed with 5 CV of buffer-B and proteins were eluted with a linear gradient from 0-100% of buffer-C (25 mM HEPES pH 7.5, 1 M NaCl, 10% glycerol (v/v), 0.5 mM TCEP) in 40 CV. Fractions containing recombinant Mediator were pooled, concentrated using a 100 kDa MWCO Amicon Ultra Centrifugal Filters (Merck) and injected onto a Superose 6 increase 10/300 GL column (GE Healthcare) pre-equilibrated in buffer-A. Fractions were analysed by SDS-PAGE and the homogenous peak fractions were pooled and concentrated to 4.5 mg/ml using a 100 kDa MWCO Amicon Ultra Centrifugal Filters (Merck). Concentrated Mediator was aliquoted, flash-frozen in liquid nitrogen and stored at −80°C until use. Pol II, TBP, TFIIA, TFIIB, TFIIE, TFIIF were purified as described^26^ and TFIIH core and the TFIIH kinase module (CAK) were purified as described^27^.

### Preparation of human Mediator-PIC complex

We formed a Mediator-PIC complex with promoter DNA at 25°C following our previously reported assembly method^7^. We used a 106 base pair (bp) Adenoviral major late promoter (AdMLP) DNA duplex (template: 5’-AGGGAGTACTCACCCCAACAGCTGGCCCTCGCAGACAGCGATGCGGAAGAGAGTGAGGACGAACGCGCCCCCACCCCCTTTTATAGCCCCCCTTCAGGAACACCCG-3’; non-template: 5’-CGGGTGTTCCTGAAGGGGGGCTATAAAAGGGGGTGGGGGCGCGTTCGTCCTCACTCTCTTCCGCATCGCTGTCTGCGAGGGCCAGCTGTTGGGGTGAGTACTCCCT-3’). Briefly, the ten subunit TFIIH was obtained by pre-incubating TFIIH core with the TFIIH CAK. Then, TBP-TFIIA-TFIIB was prebound to the AdMLP scaffold to form the upstream complex and mixed with a preformed Pol II-TFIIF complex. Following this, TFIIE and TFIIH were added in 2-minute intervals to form the PIC. Finally, Mediator was added to form the Mediator-PIC complex and this step was supplemented with ADP-BeF_3_. The mixture was incubated at 25°C for 120 min at 400 r.p.m. and aggregates were removed by centrifugation at 21,000 *g* for 5 min. The reconstituted Mediator-PIC complex was subjected to sucrose-gradient centrifugation in a 4 ml centrifugation tube (Beckmann Coulter) with simultaneous crosslinking^40^. The gradient was prepared using a BioComp Gradient Master 108 (BioComp Instruments) by mixing a 15% sucrose solution (15% sucrose, 25 mM HEPES pH 7.6, 100 mM KCl, 5 mM MgCl_2_, 5% glycerol (v/v) and 3 mM TCEP) and a 40% sucrose solution (40% sucrose, 25 mM HEPES pH 7.6, 100 mM KCl, 5 mM MgCl2, 5% glycerol (v/v) and 3 mM TCEP) supplemented with 0.25% of glutaraldehyde (v/v) crosslinker. The sample was centrifuged at 175,000*g* for 16 h at 4°C. 200 µl fractions were collected manually and the crosslinking reaction was quenched with a 10 mM aspartate and 30 mM lysine cocktail for 10 min. Fractions containing the Mediator-PIC complex were dialysed against buffer-D (25 mM HEPES pH 7.6, 100 mM KCl, 5 mM MgCl_2_, 5% glycerol and 3 mM TCEP) to remove sucrose.

### Cryo-EM data collection and processing

To prepare grids for cryo-electron microscopy (cryo-EM), Quantifoil R3.5/1 holey carbon grids were pre-coated with an amorphous carbon and glow-discharged for 45 s. 4 µl of Mediator-PIC sample was added to the carbon top and incubated for 5 min. The grids were blotted for 2.5 s and vitrified by plunging into liquid ethane with a Vitrobot Mark IV (FEI Company) set at 4°C and 100% humidity. Cryo-EM data were collected on the 300 kV FEI Titan Krios with a K3 summit direct detector (Gatan) and a GIF quantum energy filter (Gatan) operated with a slit width of 20 eV. Automated data collection was performed with SerialEM at a nominal magnification of 81,000x, corresponding to a pixel size of 1.05 Å/pixel^41^. A total of 21,894 image stacks, with each stack containing 40 frames, were collected at a defocus range of −0.3 to −5.0 µm. All movie frames were contrast transfer function (CTF) estimated, motion corrected and dose weighted using Warp^42^. Particles were picked by Warp using the trained neural network instance BoxNet, resulting in 2,309,888 particles as a starting set. Subsequent steps of image processing were performed with cryoSPARC^43^ and RELION 3.1.0^44^.

Particles were extracted with a binning factor of four and a box size of 120 pixels (a pixel size of 4.2 Å/pixel) to perform initial clean up and sorting. The first major focus of processing was to identify the best Mediator containing classes. By performing multiple rounds of heterogenous and homogenous refinements in cryoSPARC^43^, a set of 294,904 particles (Set-1) containing PIC-Mediator was identified. Set1 was re-extracted without binning and processed with RELION 3.1.0, as follows. The particles were first refined using a three-dimensional (3D) refinement followed by a focussed 3D classification with a large spherical mask (Mask-1) encompassing the region of Mediator and the Pol II stalk. This resulted in identifying the best 85,182 Mediator containing particles (Set-2). These particles were again subjected to focussed 3D refinement using Mask-1, giving rise to a Mediator-Pol II stalk reconstruction at a 4.0 Å resolution. In parallel, focussed 3D classification of Set1 with a spherical mask (Mask-2) around the TFIIH-exit DNA region helped identify the best 83,713 TFIIH containing particles (Set-3). Particles from Set-3 were then focussed 3D refined with Mask-2 to arrive at a 6.5 Å map of TFIIH (Map-2). Finally, the best Mediator-PIC containing 25,967 particles (Set-4) were identified by finding particles present in both Set-2 and Set-3. A consensus 3D refinement of Set-4 led to a final reconstruction of the entire PIC-Mediator ensemble at 4.5 Å resolution. The reported resolutions were calculated based on the gold standard Fourier shell correlation (FSC) 0.143 criterion. Post-processing of the final reconstructions of Mediator-Pol II stalk, TFIIH and PIC-Mediator were performed based on automatic B-factor determination in RELION (−85Å^2^ for Mediator-Pol II Stalk, 0 Å^2^ for TFIIH and −80 Å^2^ for PIC-Mediator) for all three final maps. Local resolution estimates were calculated using the inbuilt local resolution tool of RELION and the estimated B-factors. To assist model building, a local resolution filtered map (but unsharpened) of Mediator-Pol II Stalk was sharpened locally using PHENIX.auto_sharpen.

### Model building and refinement

Homology models of all 15 core Mediator subunits were generated by SWISS-Model^45^ using the high-resolution structures of the yeast Mediator^23^. The seven subunits of the head module and the middle module subunits MED14 and MED31 were initially built by placing the homology models into the density using rigid-body fitting in Chimera^46^. The divergent parts of these subunits were manually adjusted in Coot^47^ to fit the density. The metazoan specific subunits MED27-MED30 of the proximal tail were manually built *de novo* using Coot. The helices and strands of all subunits were placed and built with confidence in their register using bulky residues as sequence markers. Ambiguous densities of loops connecting the helices and strands were not modelled. The resulting model of the Mediator bound to the Pol II stalk was real-space refined in PHENIX^48^ and subjected to multiple rounds of manual adjustment and real space refinement in Coot and PHENIX respectively, to achieve the final model with good stereochemistry as assessed by Molprobity^49^.

The Mediator middle module was built based on the published yeast structure^23^ as follows. A conservative homology model of the middle module lacking the MED14 beam was generated from the homology models of the single Mediator subunits and placed into the Mediator-Pol II stalk focused refined cryo-EM map lowpass-filtered to 12 Å in Coot^47^. The model was stripped of flexible loops and long chain termini. We manually adjusted the trajectories of the helices in the connector and hook regions and in proximity to the knob, and parts of MED4, MED7 and MED14 that were interacting with MED31. The model was adapted to the cryo-EM density of the Mediator-PIC cryo-EM map by flexible fitting with NAMDINATOR^50^ and corrected for geometry outliers in alternating rounds of manual adjustment in Coot^47^ and real-space refinement in PHENIX^48^.

The final model was obtained by manually merging the resulting Mediator-Pol II stalk model with the atomic model of the PIC^26^ lacking the Pol II stalk. The Mediator-PIC model was first manually examined against the Mediator-PIC overall map in Coot and then real space refinement was performed with Phenix. The final model showed good stereochemistry as assessed by Molprobity^49^ (Extended Data Table 1).

### Structure base sequence alignment

Structure based sequence alignment comparing the respective Mediator subunits from humans and *Schizosaccharomyces pombe* was performed using the Expresso algorithm of the T-Coffee server^51^. The alignments were manually corrected for discrepancies and the secondary structure labels were assigned using the program ESPript from the ENDscript server^52^.

## EXTENDED DATA FIGURES

**Extended Data Figure 1.**
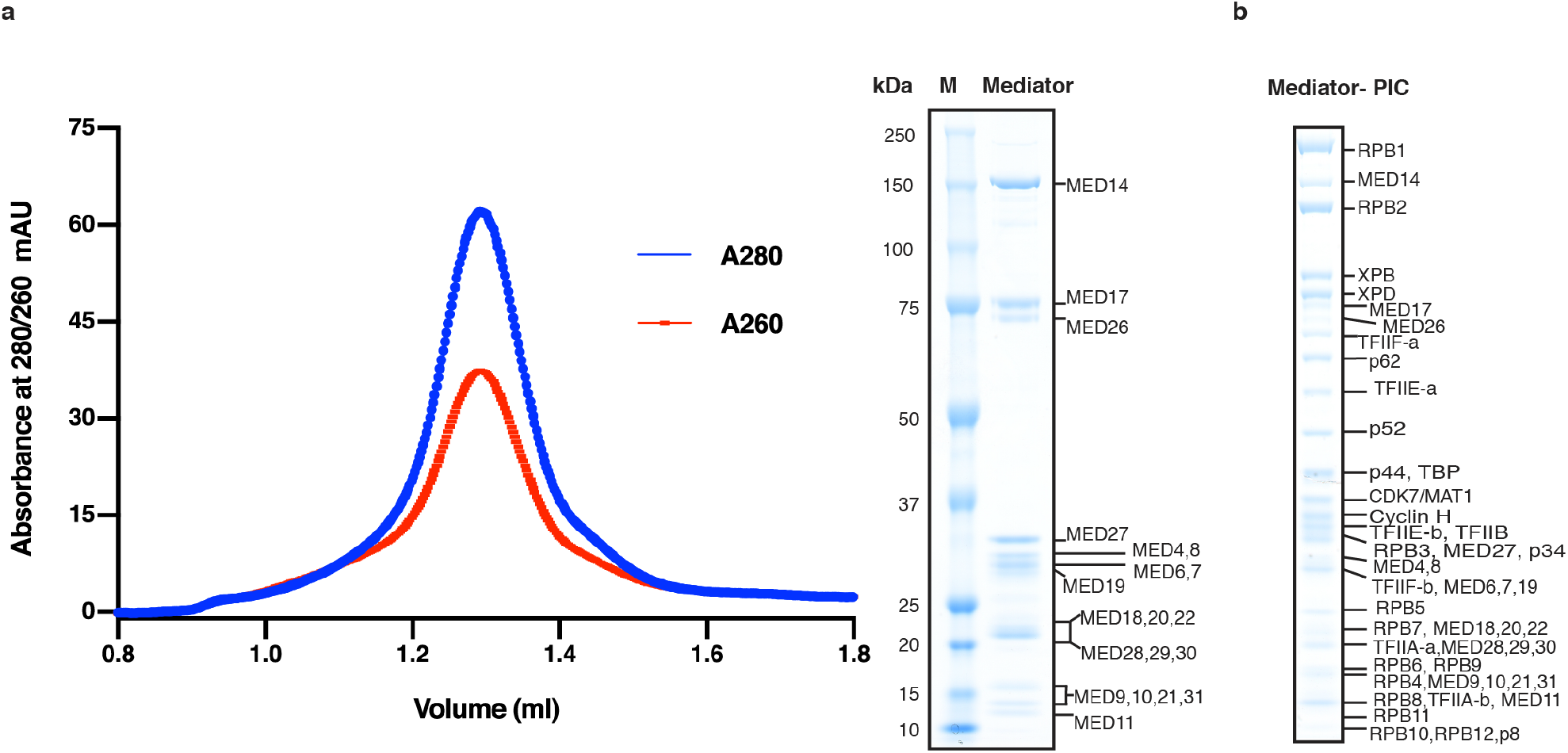
Preparation of the human Mediator-PIC complex. **a.** Size exclusion chromatography of recombinant human Mediator shows a single peak. SDS-PAGE analysis shows the presence of 20 Mediator subunits, confirmed by mass spectrometry (not shown). **b.** SDS-PAGE analysis of human Mediator-PIC complex after isolation from a sucrose gradient.

**Extended Data Figure 2.**
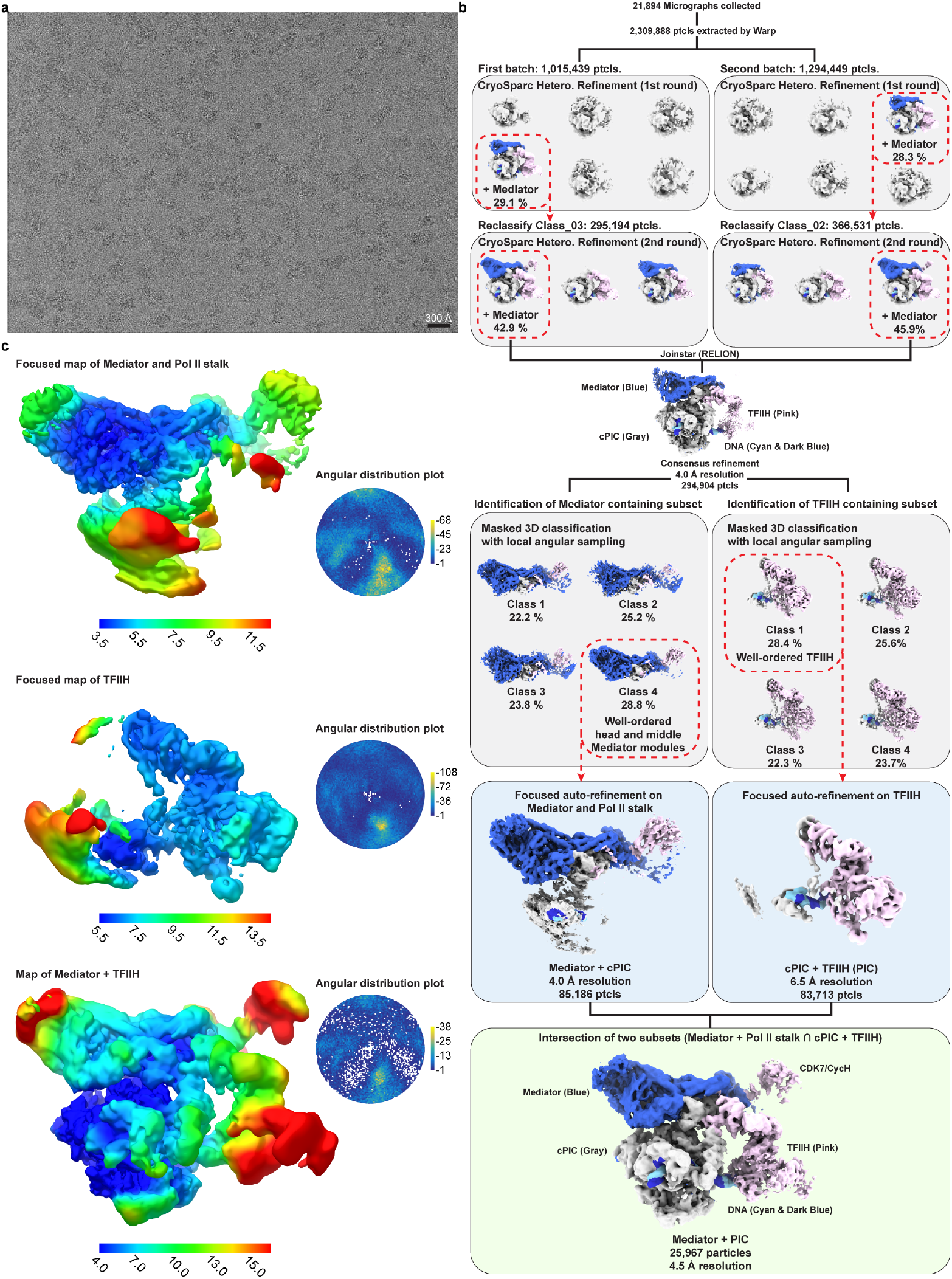
Cryo-EM data processing. **a.** Representative cryo-EM micrograph of human Mediator-PIC complex. Scale bar represents 300 Å. **b.** Processing tree describing particle classification. Reconstructions that gave rise to maps used for model building are indicated (blue for focused maps, green for overall maps). Regions corresponding to cPIC are colored gray, TFIIH (including the CAK) are colored pink, DNA is colored dark blue and cyan, and Mediator is colored royal blue. **c.** Reconstructions colored by their local resolution as estimated using RELION. In the angular distribution plots color indicates particle representation (white areas indicate unpopulated angles).

**Extended Data Figure 3.**
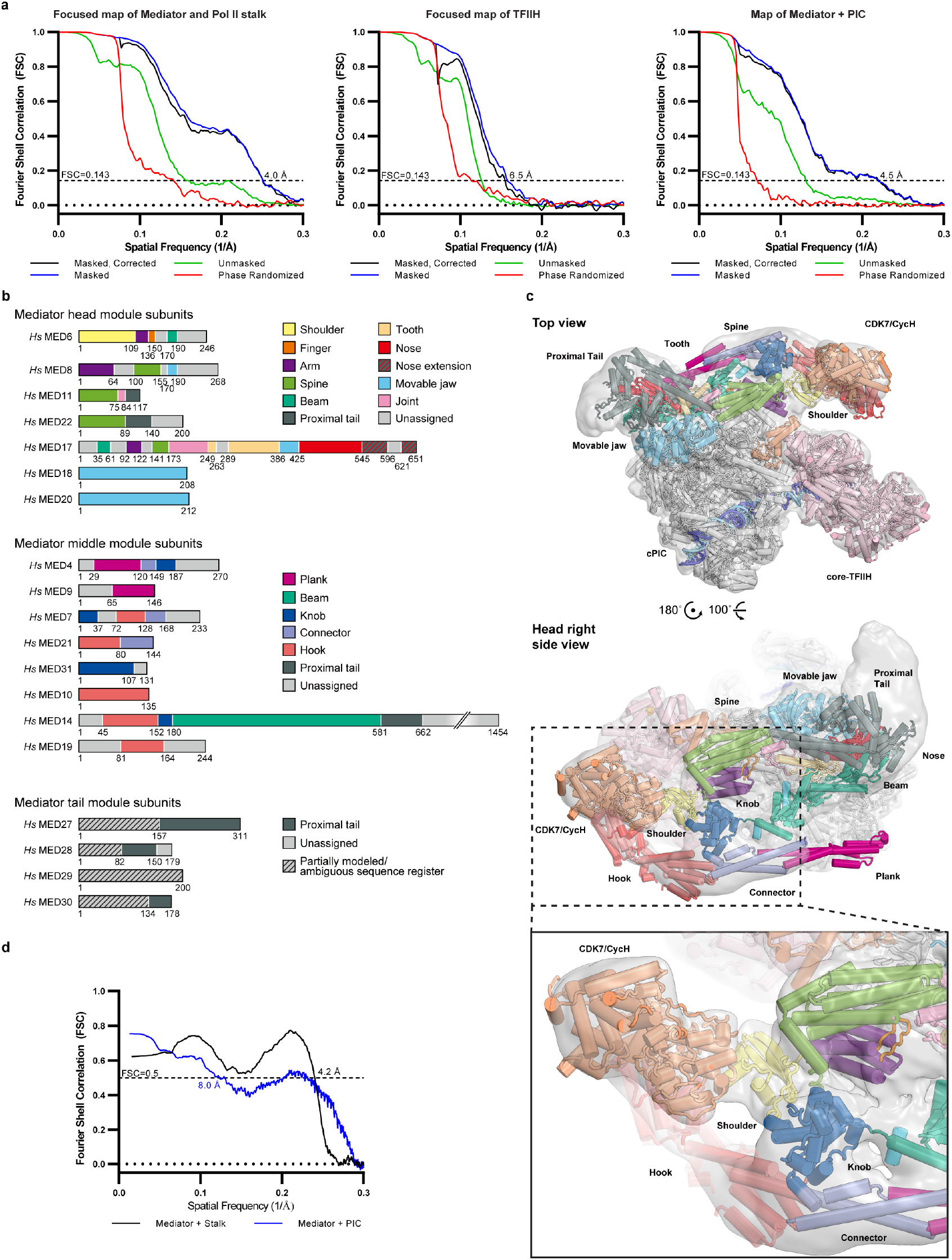
FSCs of reconstructions and cryo-EM density. **a.** Solvent-corrected ‘gold-standard’ FSCs for the reconstructions shown in Extended Data Fig. 2, panel c. Unmasked (green), masked (blue), and phase-randomized (red) FSCs are also shown. **b.** Schematic representation of Mediator subunit domain architecture. Regions contributing to submodules are coloured according to the *S. pombe* cMed structure^23^. Unassigned regions are coloured grey. **c.** Local resolution-filtered map of Mediator+PIC with the fitted structure. Inset shows zoomed in view to the CAK fitting into our density. **d.** Model-to-map FSCs, showing in blue the fit of the overall structure to the Mediator + PIC and in black the fit of the Mediator-Pol II stalk model to their corresponding maps.

**Extended Data Figure 4.**
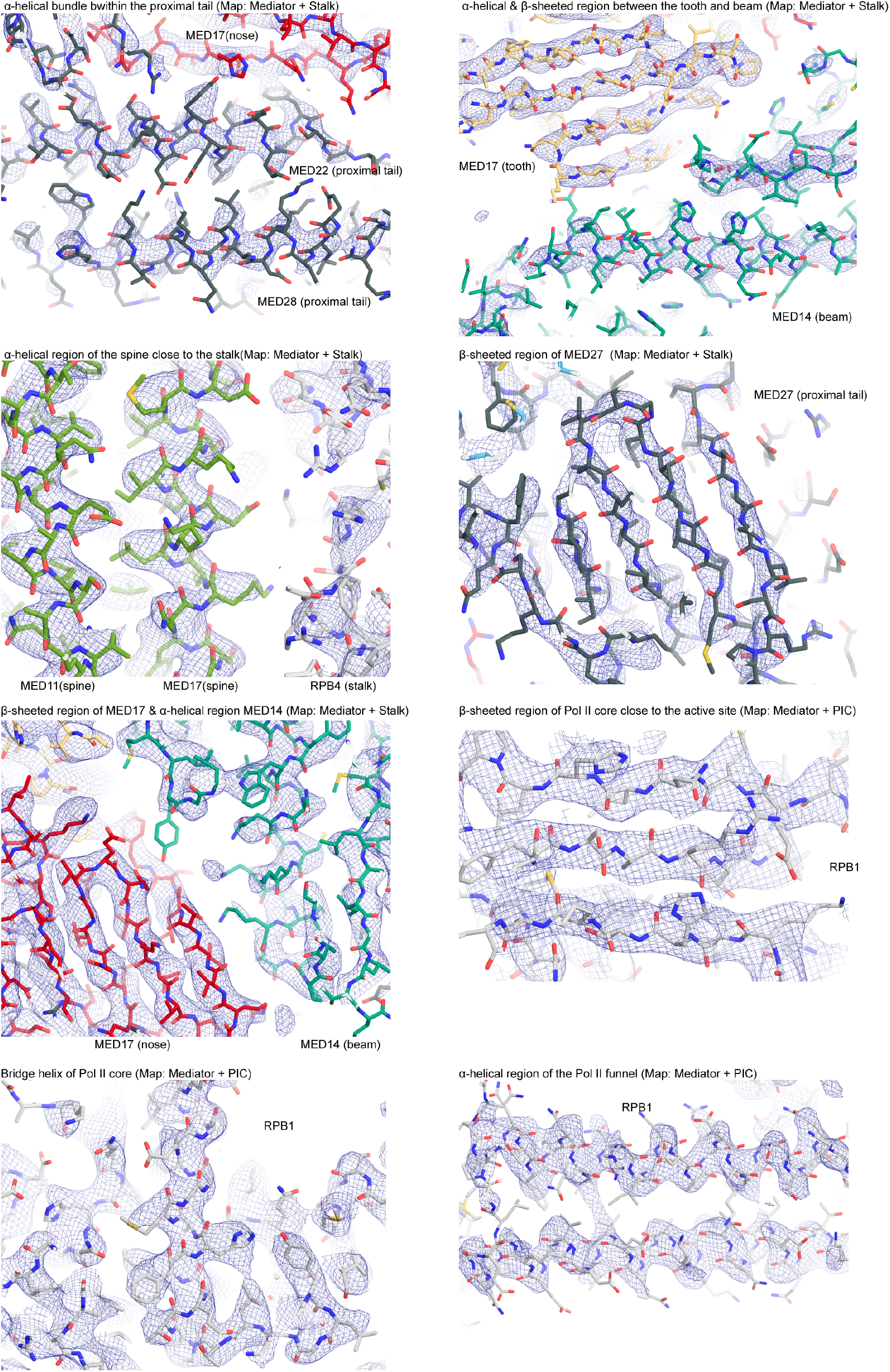
Quality of cryo-EM densities. Sections of focussed-refined Mediator-stalk cryo-EM density overlaid with their respective atomic models. Densities are illustrated as a blue mesh, and sticks are shown for the model colored as in Extended Fig. 3, panel c.

**Extended Data Figure 5.**
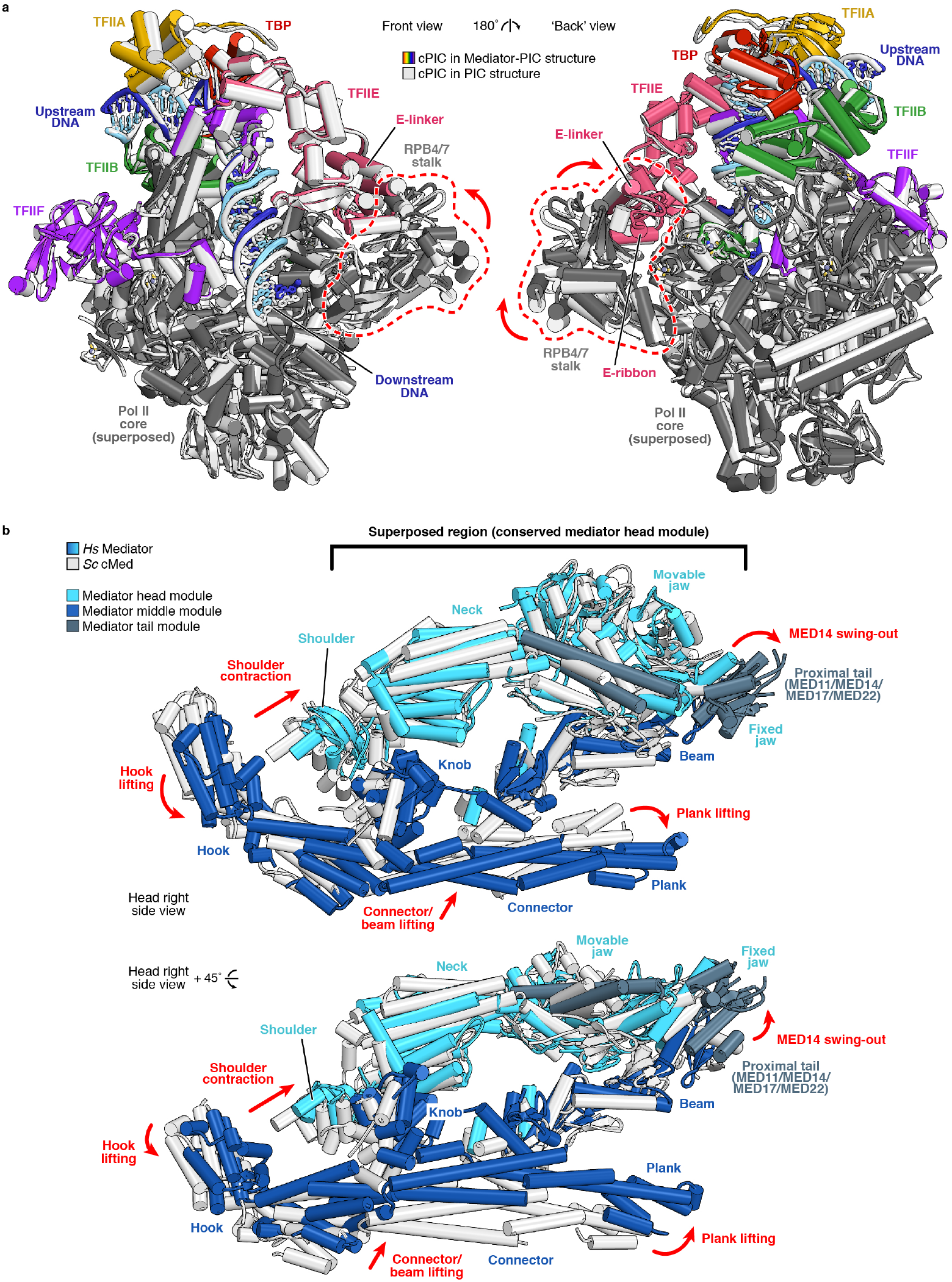
Additional structural comparisons. **a.** Comparison of the human Mediator-PIC structure (this study) with the free PIC structure^26^reveals a different orientation of Pol II RPB4-RPB7 stalk. The core PIC regions of the Mediator-PIC (in colour) and free PIC (in white) structures were superposed on the 10-subunit Pol II core. Mobile elements in the stalk are indicated and conformational changes between complexes are depicted by red arrows. **b.** Comparison of the yeast and human Mediator-PIC structures reveals a different relative orientation of the Mediator middle module. The human (in color) and yeast^9^ (in white) Mediator-PIC structures were superposed on the well-conserved neck and fixed jaw domains of the Mediator head module. Conformational changes of Mediator submodules are depicted by red arrows. The proximal tail region of human Mediator was omitted for clarity.

## EXTENDED DATA TABLES

**Extended Data Table 1.** Cryo-EM data collection and processing information.

| Sample | Mediator-PIC |  |  |
| --- | --- | --- | --- |
| Subset | Mediator-PIC | Mediator-Pol II stalk | TFIIH |
| **Data acquisition** |  |  |  |
| Voltage |  | 300kV |  |
| Magnification |  | 81,000 |  |
| Pixel size |  | 1.05 Å |  |
| Electron dose |  | 45 e<sup>-</sup>/Å<sup>2</sup> |  |
| Frame number |  | 40 |  |
| Defocus range |  | 0.3-5.0 |  |
| **Data Processing** |  |  |  |
| Final number of particles | 25,967 | 85,182 | 83,713 |
| Map resolution at 0.143 FSC | 4.5 Å | 4.0 Å | 6.5 Å |
| Local resolution range | 4.0 – 15.0 Å | 3.5 – 11.5 Å | 5.5 – 13.5 Å |
| Map sharpening B-factor | -80 Å<sup>2</sup> | -80 Å<sup>2</sup> | 0 Å<sup>2</sup> |
| **Model composition** |  |  |  |
| No. of atoms | 96,178 |  |  |
| Protein | 11,641 |  |  |
| DNA | 136 |  |  |
| Ligand | Zn: 18<br>ADP: 1<br>Mg: 3<br>SF4: 1<br>BEF: 1<br>UNK:24 |  |  |
| **Model validation** |  |  |  |
| MolProbity score | 1.72 |  |  |
| Clash score | 8.16 |  |  |
| Rotamer outliers (%) | 0.00 |  |  |
| Cβ outliers | 0.02 |  |  |
| **Ramachandran plot** |  |  |  |
| Favoured | 96.03 |  |  |
| Allowed | 3.95 |  |  |
| Outliers | 0.02 |  |  |

## SUPPLEMENTARY INFORMATION

**Supplementary Figure 1. | Structure-based alignment of Mediator subunit sequences.** These primary sequence alignments were obtained by manual comparison of our PIC-bound human Mediator structure with the structure of the *S. pombe* core Mediator. Secondary structure elements observed for human (top) and *S. pombe* subunits (bottom) are indicated.

## MED6

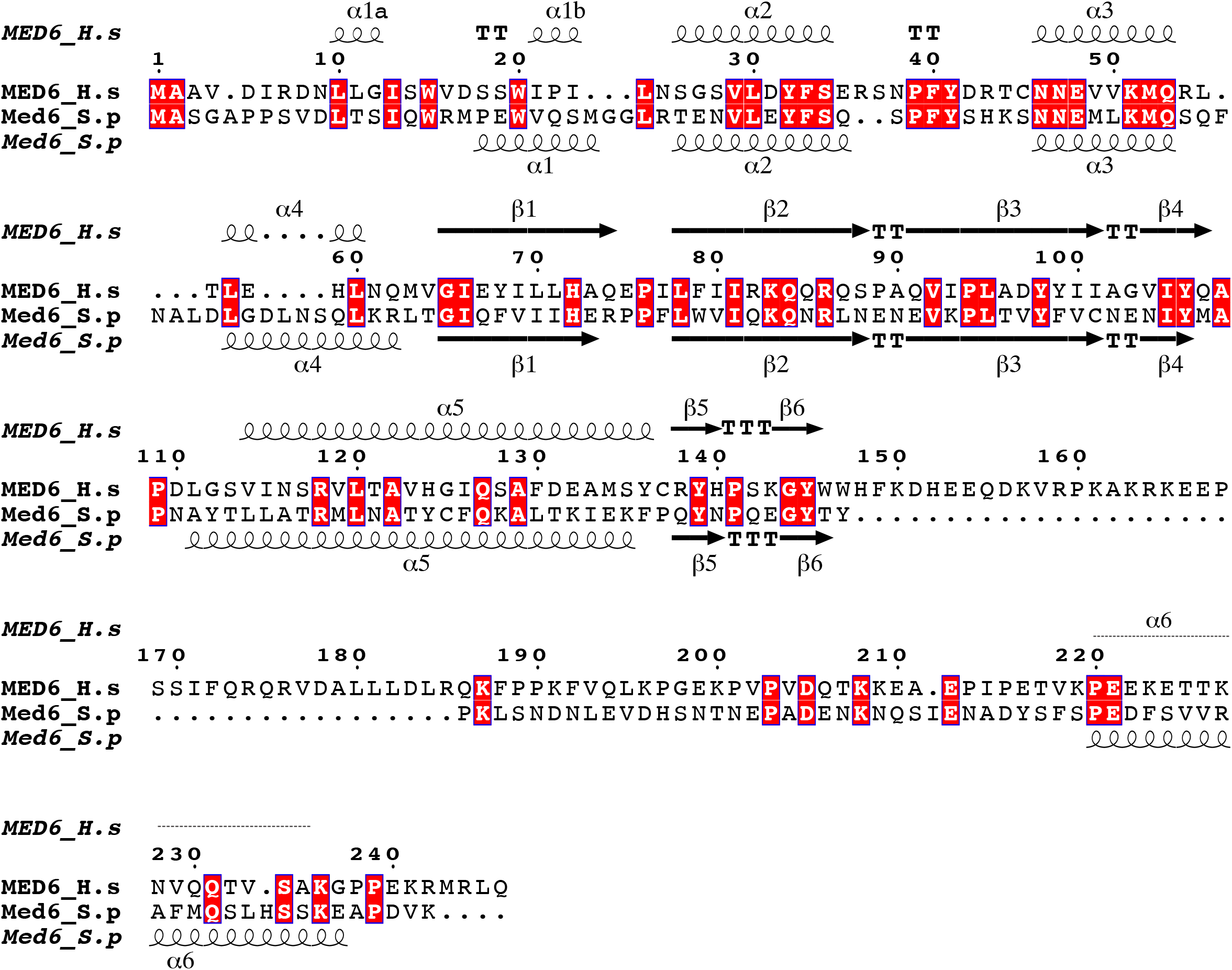

## MED11

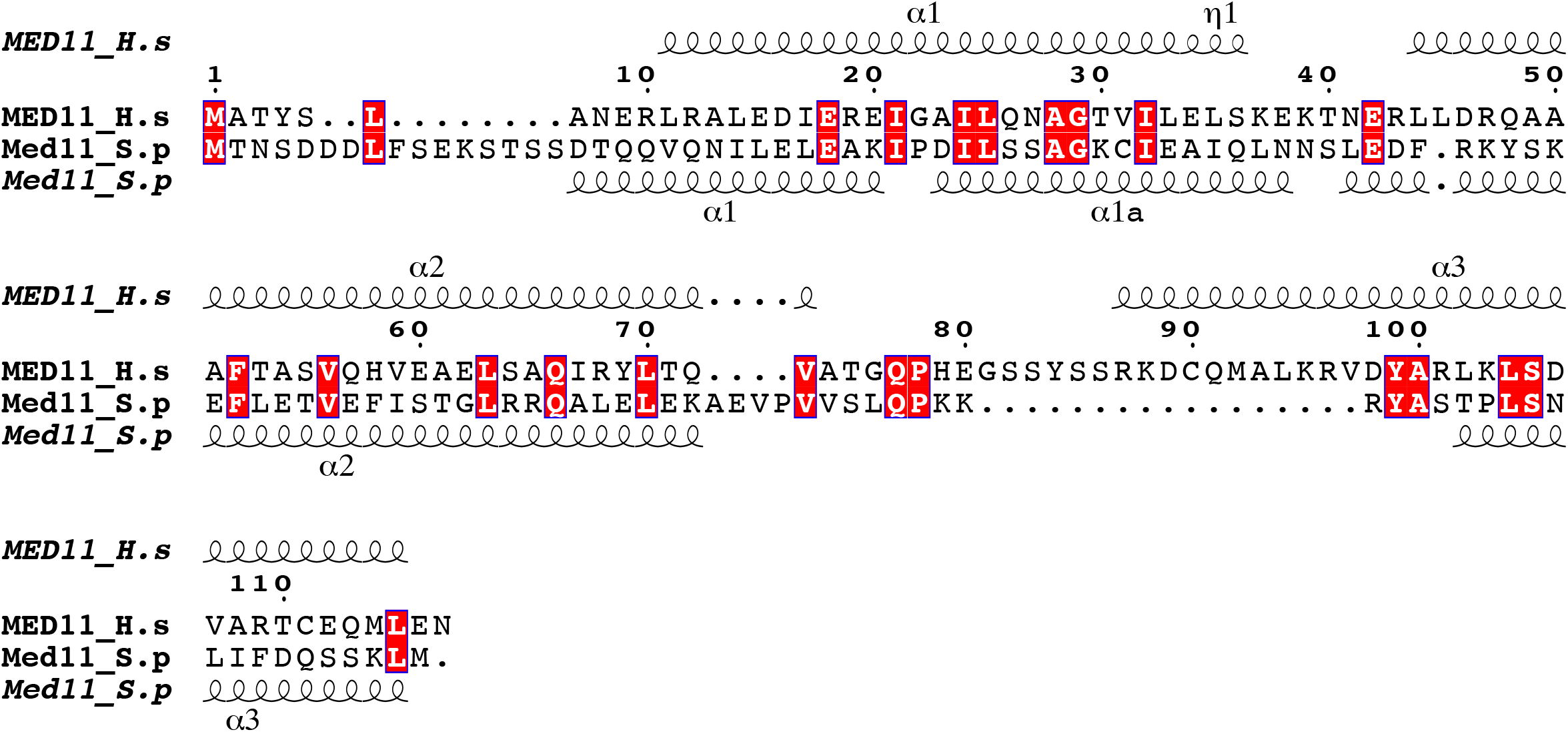

## MED8

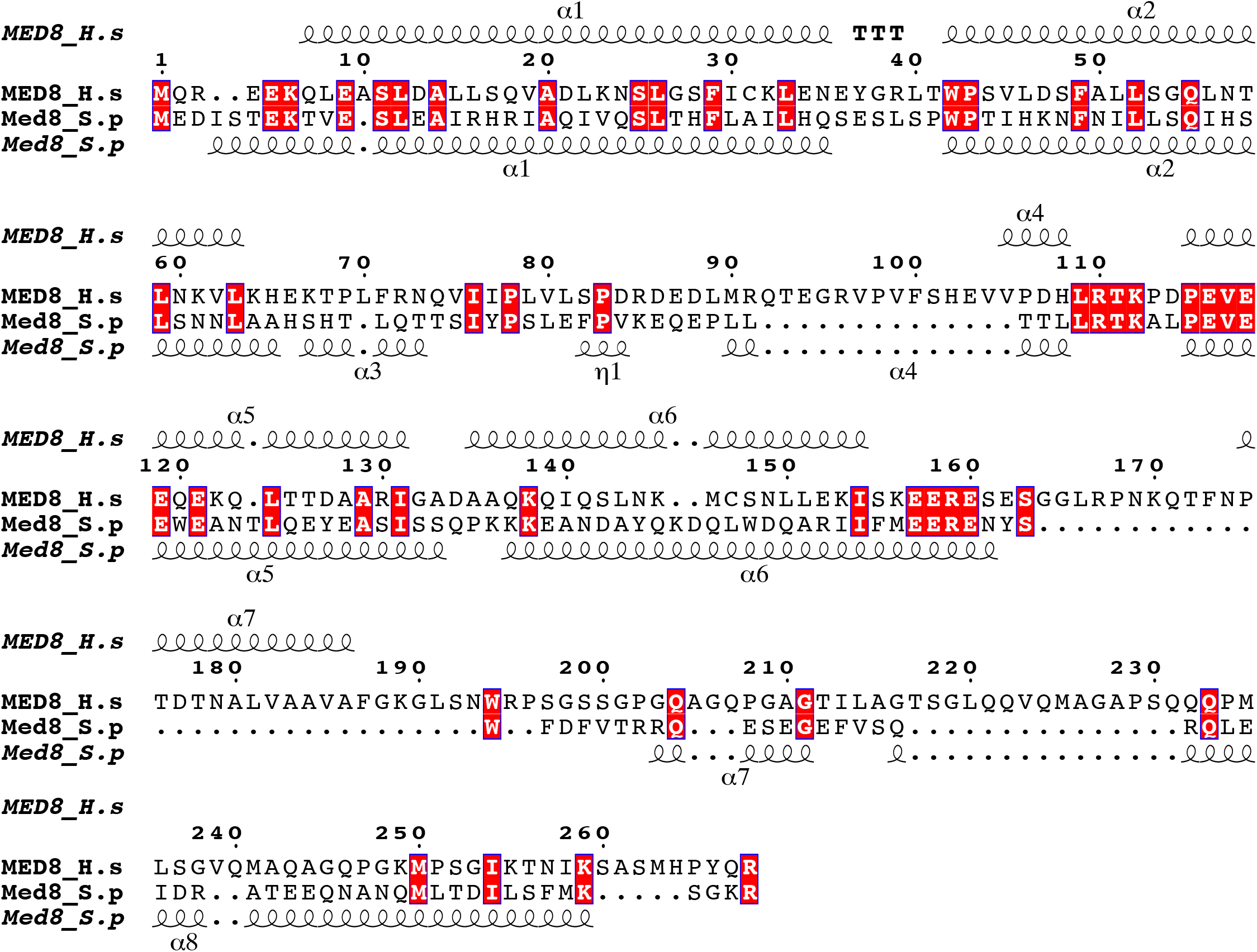

## MED22

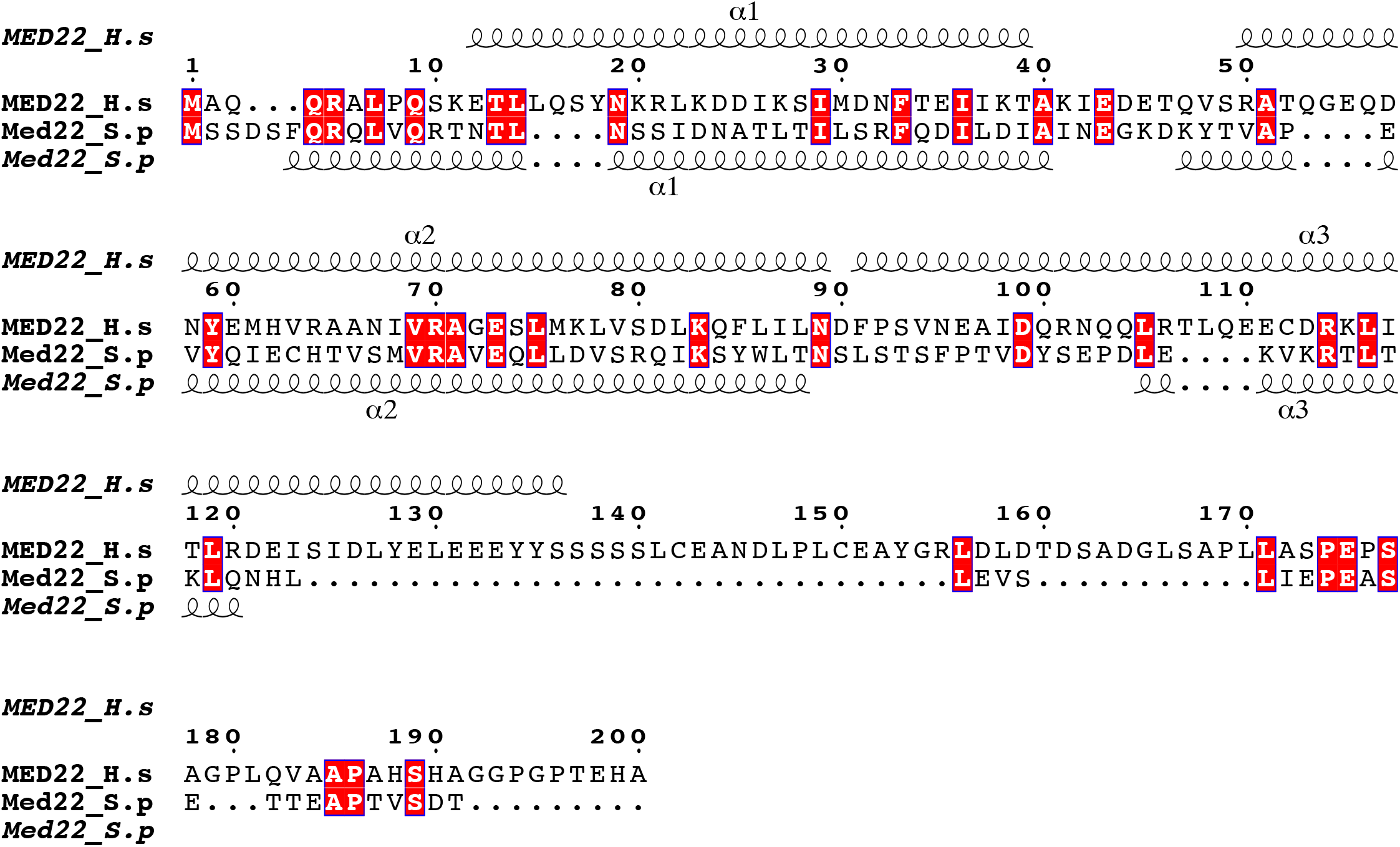

## MED18

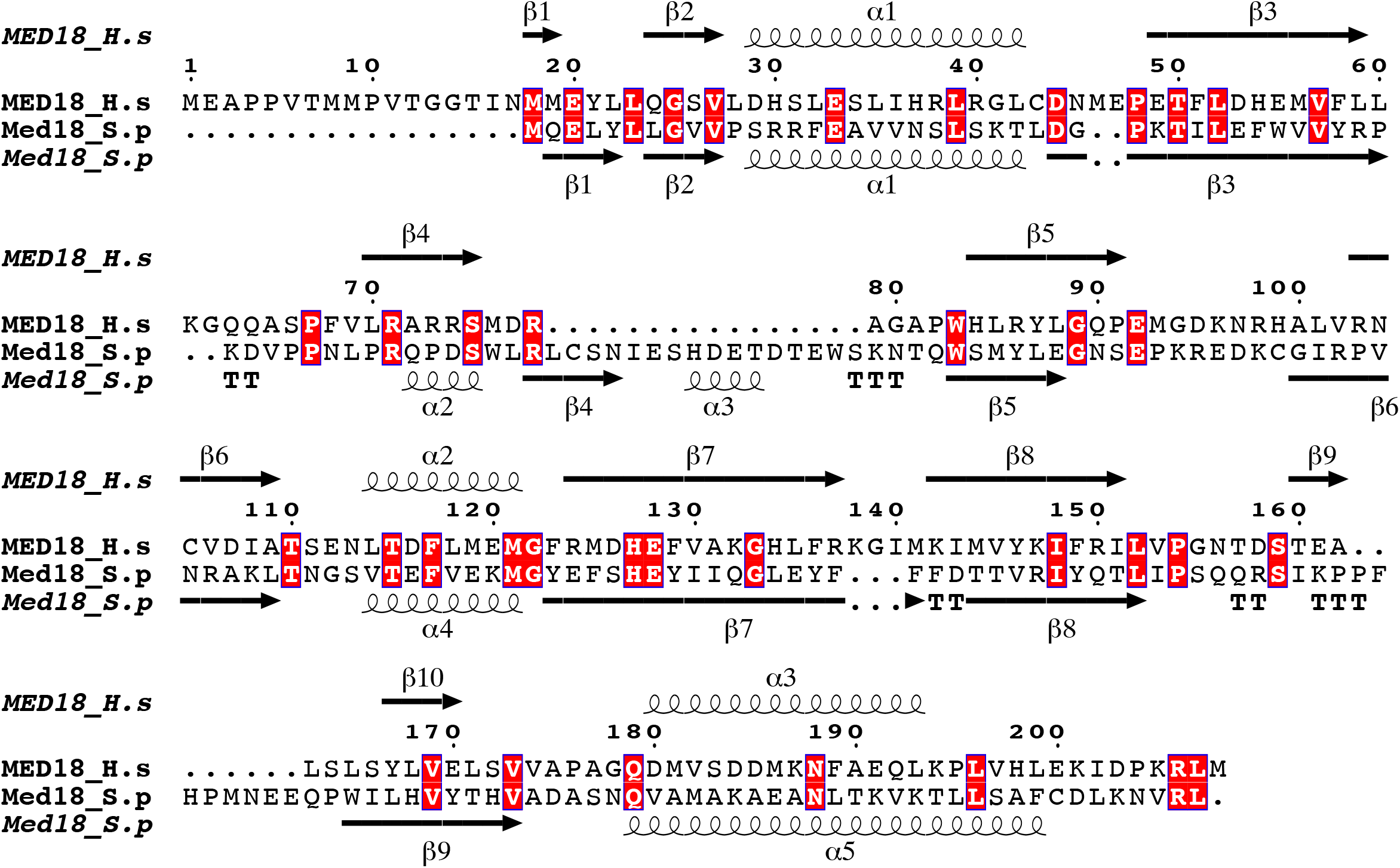

## MED20

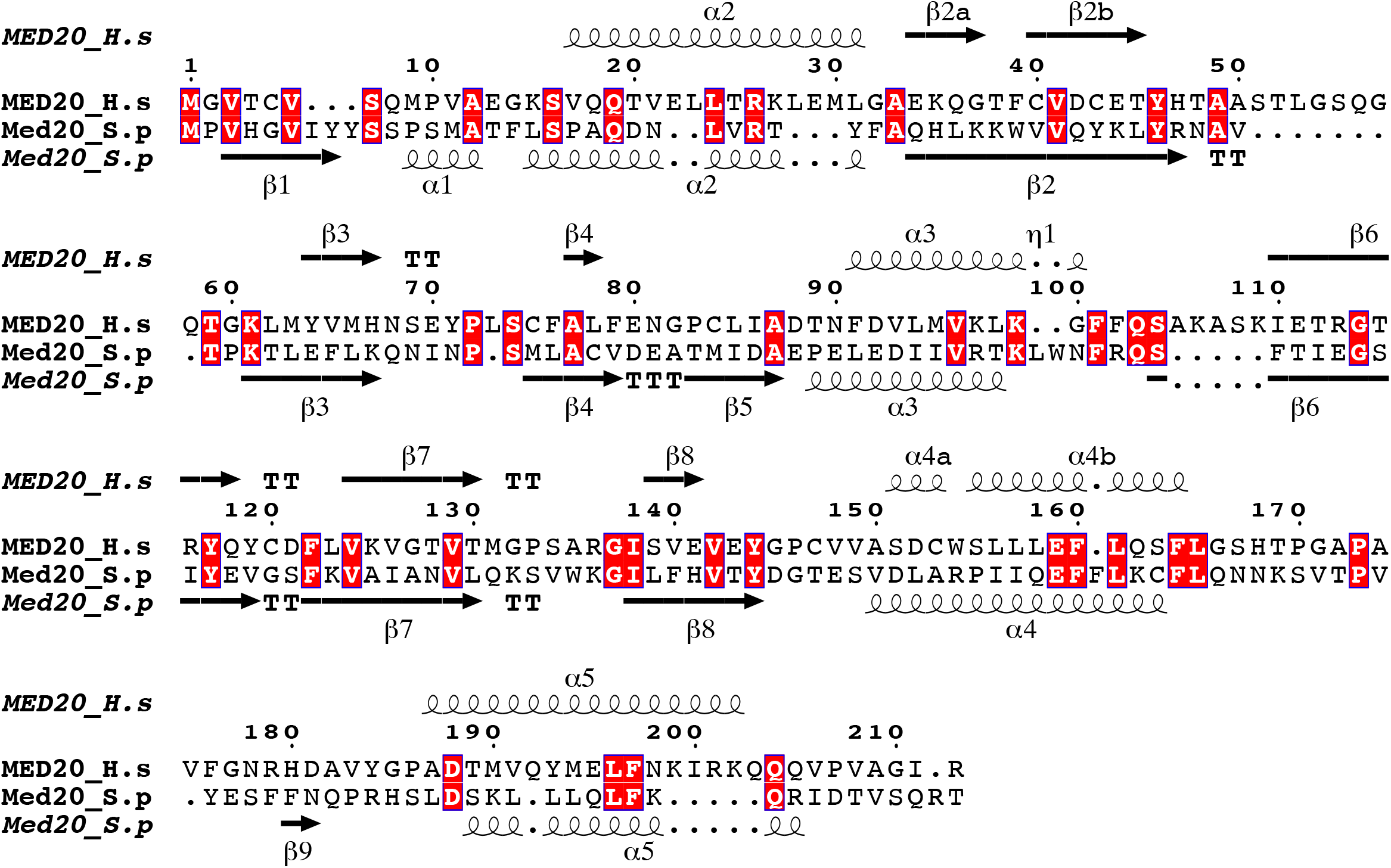

## MED17

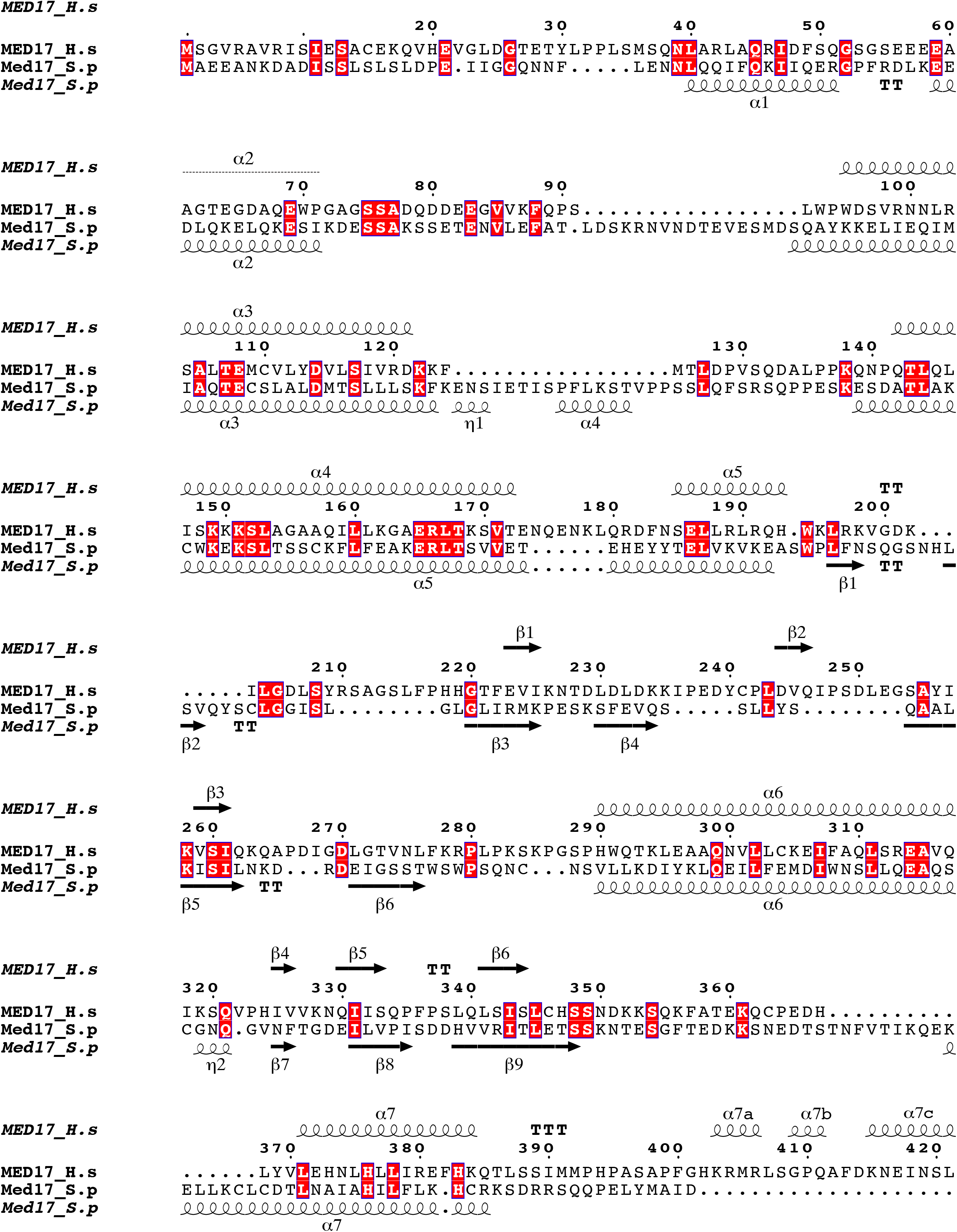
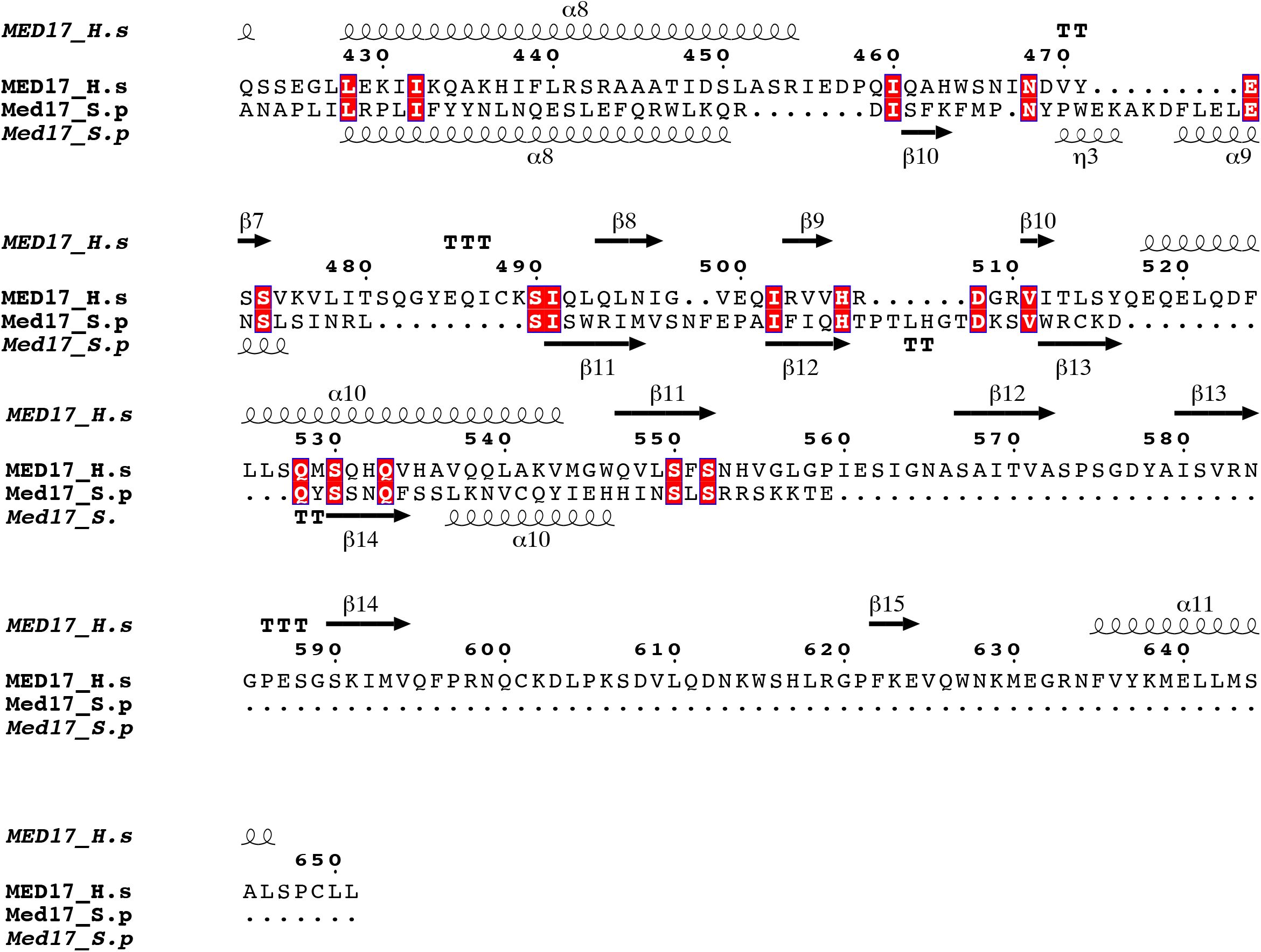

## MED14

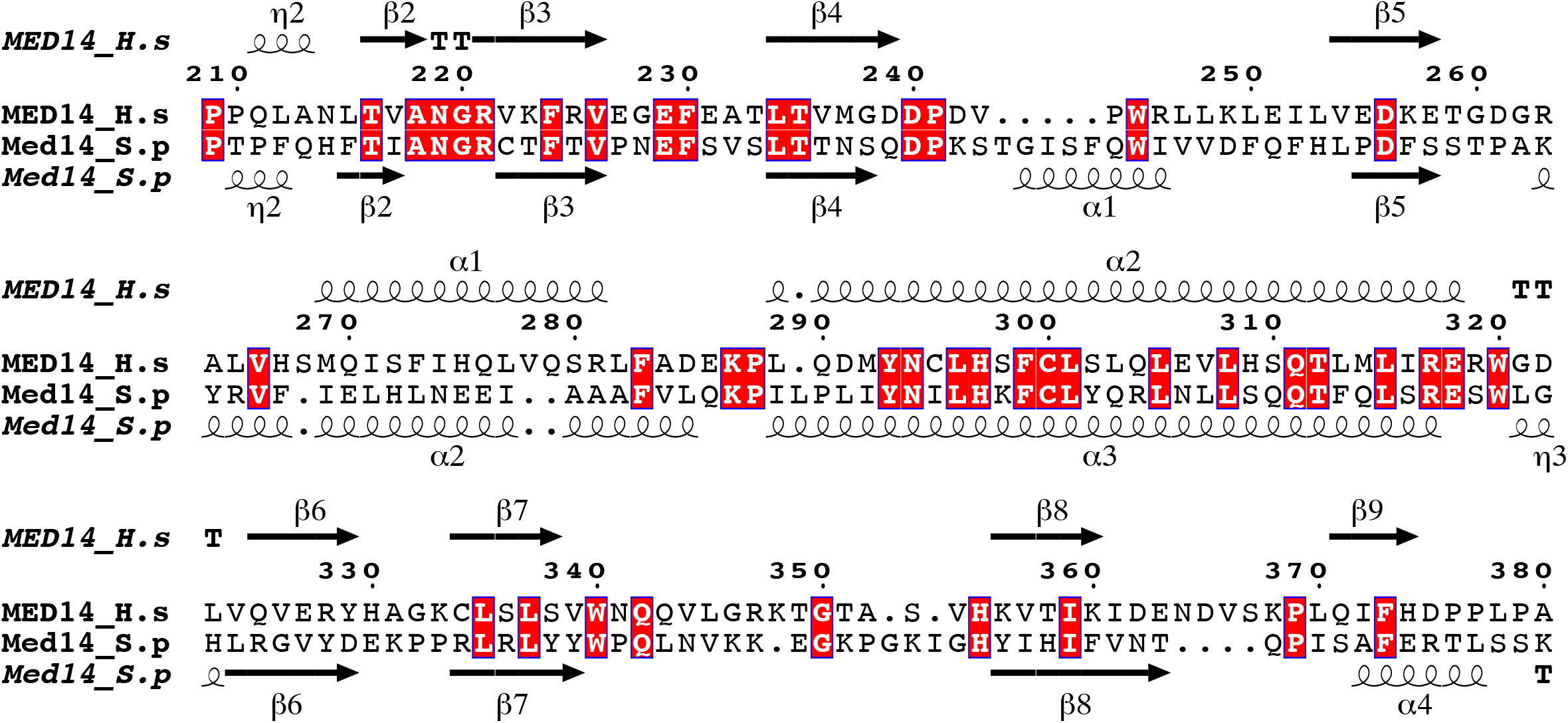
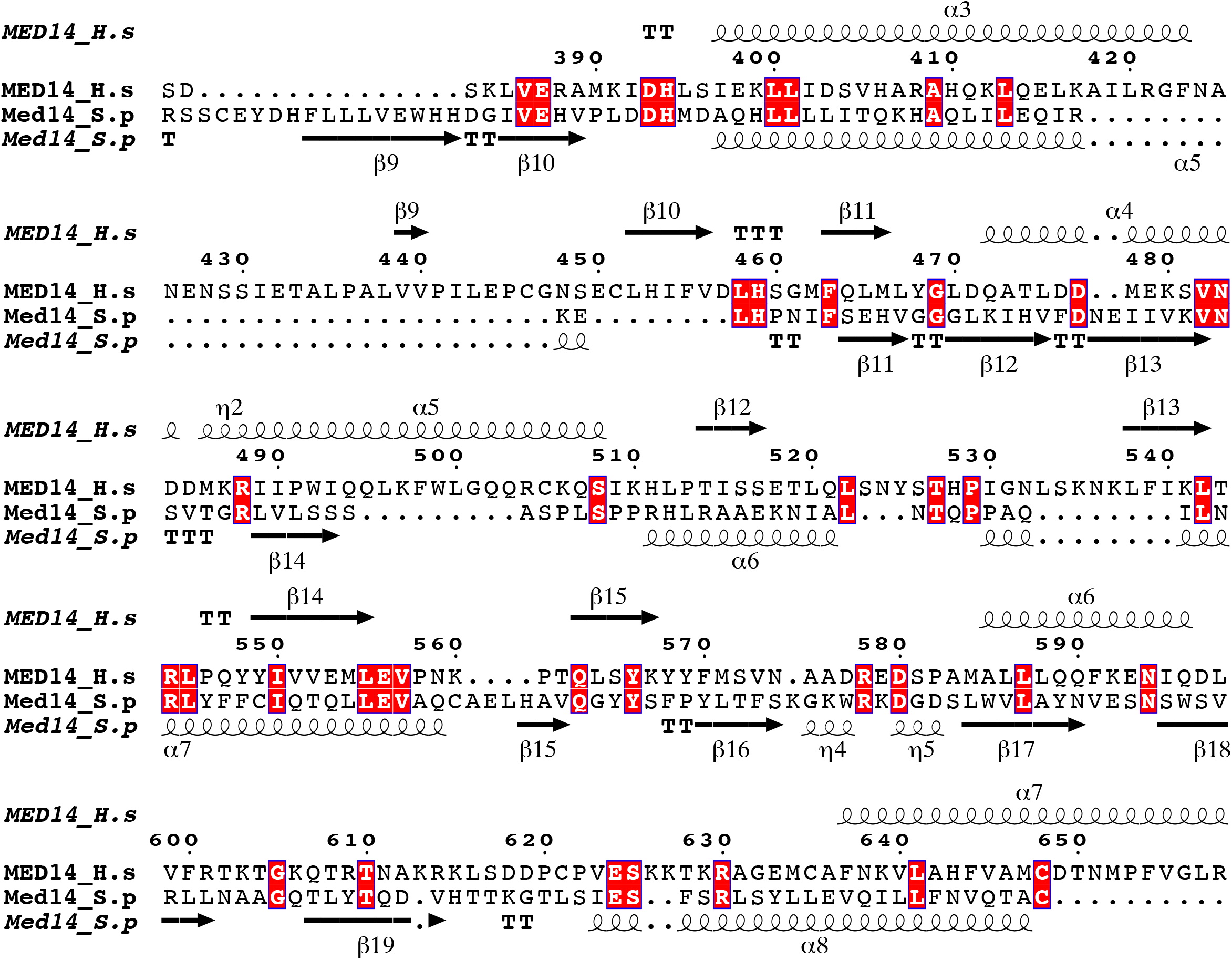

## Notes

### Competing Interest Statement

The authors have declared no competing interest.

